# League of Brazilian Bioinformatics: a competition framework to promote scientific training

**DOI:** 10.1101/2020.12.17.423357

**Authors:** L. M. Carvalho, N. A. R. Coimbra, M. R. C. Neves, N. J. Fonseca, M. A. Costa, E. C. A. Horacio, R. Riyuzo, F. F. Aburjaile, S. T. Nagamatsu

## Abstract

**Background:** the scientific training to become a bioinformatician includes multidisciplinary abilities, which increase the challenges to professional development.

**Competition framework:** in order to improve and promote the ongoing training of the Brazilian bioinformatics community, we organize a national competition, with the main goal to develop human resources and abilities in Computational Biology at the national level. The competition framework was designed in three phases: 1) a one-day challenge composed of 60 multiple-choice questions covering Biology, Computer Science, and Bioinformatics knowledge; 2) five Computational Biology challenges to be solved in three days; and 3) development of an original project evaluated during the 15th X-meeting.

**Results:** the first edition of the League of Brazilian Bioinformatics (LBB) counted 168 competitors and 59 groups, distributed into undergraduate students (14.4%), graduate students (12.6% master and 16.8%, Ph.D.), and other professional fields. The first phase selected 46 teams to proceed in the competition, while the second phase selected the three top-performing teams.

**Conclusion:** during the competition, we were able to stimulate teamwork in the main areas of Bioinformatics, with the engagement of all research-level competitors. Furthermore, we identified opportunities to deliver and offer better training to the community and we intend to apply the acquired experience in the second edition of the LBB, which will occur in 2021.

**Supplementary information:** Supplementary data are available at *Bioinformatics*

## 1 Introduction

Bioinformatics is a multidisciplinary field that encompasses mainly biology, mathematics, statistics, chemistry, physics, and computing. This integrative characteristic turns it into an area with great challenges, including development and opportunities for learning and training. Furthermore, the emergence of highly skilled professionals in multiple fields has also been growing due to the increased demand for interdisciplinary areas. Still, despite the development of the Bioinformatics field, there is a lack of educational materials in developing countries. The main challenge is associated with the difficulty in integrating bioinformatics with the various departments that demand this knowledge (Kandemir-Cavas *et al.*, 2018).

Encouraging scientific training to the next generation of researchers and transfer knowledge in Bioinformatics is essential to students and young professionals, especially at the beginning of their careers (Wang *et al.*, 2018). It can be improved by formal training, university subjects, and using computer tools, such as educational materials in applied courses, undergraduate projects for students in bioscience departments, and also hackathons (Nunes *et al.*, 2015; Militello, 2013; Kandemir-Cavas *et al.*, 2018; Brown, 2016; Weisman, 2010; Vyahhi *et al.*, 2012). Moreover, the up-to-date term BioHackathon came up to refer to hackathons that aim to explore relevant topics in the biomedical and life sciences field, and several have already been carried out (Garcia *et al.*, 2020). In the need for innovative ways to evaluate and improve bioinformatics training, we highlight the important role of hackathons and competitions as methods to foster academics and professionals training in Computational Biology.

Indeed, connecting students to problem solving challenges has been an approach that has demonstrated such an improvement in the ability to conduct research while expanding their opportunities to be absorbed by different sectors of industry and market (Pathanasethpong *et al.*, 2019; Mason et al., 2009). With the gradual increase of professionals in Bioinformatics and the specialization that is requested, competitions for knowledge in this area began to arise (Kienzler and Fontanesi, 2017; Lawson et al., 2020; Connor et al., 2019; Pathanasethpong et al., 2019; Wang *et al.*, 2018; Silver *et al.*, 2016). Furthermore, hackathons have been applied in specific areas to explore public data, increase educational and innovation opportunities, create biological insights (2019 Model “Metrics” Challenge, Health Hackathons and NCBI’s Virus Discovery Hackathon), and even solve new challenges and improve the state of the art (AlQuraishi, 2019; Zhou et al., 2019; Lawson et al., 2020). The experiment showed an effective development and reproducibility of the analyses performed during the competition by the teams. Moreover, Wang et al. highlighted the educational value of hackathons, which are necessary to develop a standardized assessment of knowledge and evaluate the larger sample size of the surveyed participants.

According to the Web of Science, in the Brazilian context the number of publications in Bioinformatics by national authors is increasing, impacting the number of citations peer-reviewed in the field, even after progressive series of cuts in the governmental funding agencies (Analytics, 2017). We also found initiatives in the country to improve education in Bioinformatics as a manual for professors, who want to implement initial training for undergraduate or graduate students in life sciences (Mariano et al., 2019).

Taking into account all these aspects mentioned, the ISCB Regional Student Group Brazil (RSG-Brazil) assembled the League of Brazilian Bioinformatics (LBB). LBB is a national competition in Bioinformatics that aims to train, stimulate competitions, and promote challenges to integrating the Bioinformatics and Computational Biology community. In this article, we presented the experiences of the LBB held in 2019 as a positive experience, contributing to an increase in training, students’ engagement, practical bioinformatics skills, and knowledge transfer.

## 2 Approach

### 2.1 League of Brazilian Bioinformatics (LBB)

The LBB (https://lbb.ime.usp.br/home) is a national biennial competition, which had its first edition in 2019. The main goal of LBB was to test the abilities and competencies of students and Brazilian bioinformaticians in the broad areas of Computational Biology. It was also an encouraging action to support, attract, and discover talents nationally. The LBB was assembled by the RSG-Brazil (http://rsg-brazil.iscbsc.org/) in association with the Brazilian Association for Bioinformatics and Computational Biology (AB3C: http://www.ab3c.org). LBB was conceived to achieve specific objectives: (i) Encourage the continuous training of human resources in Bioinformatics through participation in competitions; (ii) Train human resources in the production of events in Bioinformatics; (iii) Stimulate and promote the organization of future bioinformatics competitions both nationally and internationally; (iv) Promote the integration of the Bioinformatics and Computational Biology community in the country and collaboration between LBB participants.

### 2.2 The competition framework

The competition followed a well-defined statute that contained all the rules to the three phases and the competencies expected for the team’s assembling (Figure 1). Each team was composed of two or three competitors including a maximum of one Ph.D. per group. Also, professors and associate researchers were not allowed as team members. The competition framework was designed in three phases, running through a four-month competition. The complete regulation of LBB 2019 and its phases are depicted in the supplementary material.

**Fig. 1.**
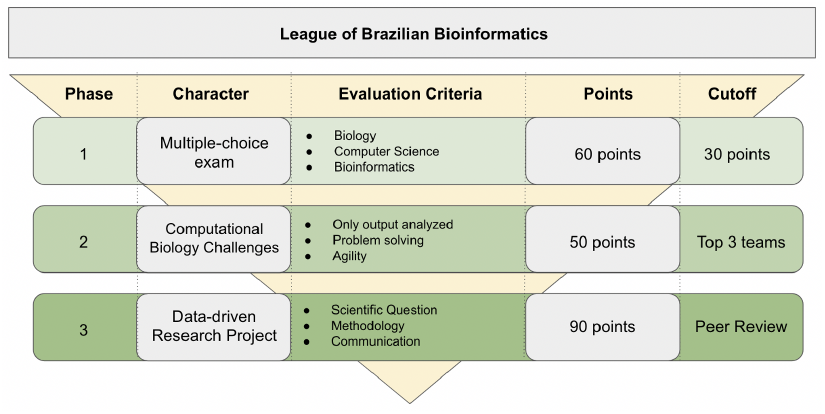
League of Brazilian Bioinformatics (LBB) framework.

First phase: One exam was composed of 60 multiple-choice questions, granting one point for each correct answer, encompassing three areas: Computer Science, Biology, and Bioinformatics. It was held online on August 04th and it lasted 5 hours and 3 minutes, alluding to the synthesis of DNA strands that have a 5 ‘->3’ sense. To achieve the second phase, the teams were required to complete at least 50% of the correct answers in the exam (cutoff 30 points). The test was performed in Google Forms and the results were analyzed automatically.

Second phase: Programming and problem-solving skills, disposed of five computational biology challenges available to competitors for 3 (three) days. Each challenge included different sub-questions totalling 10 (ten) points in the final grade. The scores were computed and the top three teams were accepted in the third phase of the LBB. The test started at 8 AM (BRT) on September 13th (Friday) and remained available for resolution until 11:59 PM (BRT) on September 15th (Sunday). The second phase test was implemented and automatically corrected in the Stepik platform (https://welcome.stepik.org/en).

Third phase: Development of an original data-driven research project with public data-sets that were presented by the teams at the X-meeting 2019 conference (https://www.x-meeting.com/events/home) on November 01st. The teams delivered a written project a week before the finals to be analyzed by three invited evaluators. The project presentations were made in the form of 15-minute seminars, explaining the scientific question and the methodologies used, as well as the results obtained. The evaluation criteria observed encompassed: (i) the existence of a well-defined scientific question, (ii) the appropriate choice of methodologies to answer that question, (iii) the exploration and proper interpretation of results obtained, (iv) the clarity and creativity of the presentation of the project, and (v) the social and/or environmental impact. The grades of the third phase had a fixed score of five criteria, judges could score 0 to 5 points on each question. The referees were selected based on previous experience on the topics of Biological Sciences, Computer Science, and entrepreneurship. If the judging panel deemed it necessary to add a bonus to the group, reaching a maximum of 90 points to the team score. Judgment criteria were based on content presentation: (a) scientific question; (b) choice of methodology; (c) discussion and results, (d) project presentation, and (e) bonus.

### 2.3 LBB Match: improving networking to team building

Using matching apps as a model, we collected personal background in Bioinformatics, geographic information, and team members’ preferences, which included bioinformatics background, geographic location, level of education, and programming level. Accounting for the multidisciplinarity character ofBioinformatics participants were placed into groups considering their primary discipline in higher education, according to their preferences. Team builder also had as a premise that participants with a complete Ph.D. were never placed in the same team, and at least one integrant would be enrolled in a higher education institution, satisfying the competition’s rules.

After team building, all participants were contacted by email to accept or reject the members.

## 3 Organizing committee

The organizing committee was composed of nine members distributed into three categories: (i) social media and customer support, (ii) legal and financial, and (iii) board exam.

## 4 Analyses strategy

### 4.1 Discrimination of multiple-choice items

Item Response Theory (IRT) is a methodology applied in assessments of different areas to describe the relationship between the level of the latent trait (ability-θ) with the characteristics of the observed items, and the person’s (or group’s) responses to each item (Yang *et al.*, 2014). A reasonable assumption is that each participant who responds to a test item has some amount of associated ability. Thus, each participant can be associated with a numerical value (score), which fits it on the ability scale. This ability score will be indicated by the Greek letter theta (θ). In the first phase test of the LBB all groups were classified according to their ability level with an associated-score ranging from 0 to 60. This score was denoted by T(θ). We could also determine how the probability of a participant with a certain ability θ would provide a correct answer to the item. This probability was denoted by P(θ). In the case of the typical test item, this probability should be small for low-capacity test takers and large for high-capacity test takers (Baker, 2001).

The Two Parameter Logistics (2PL) model allows estimating the probability of someone answering an item (of difficulty, b) correctly. Furthermore, it also allows estimating the discrimination of item (a), which is the ability of an item to distinguish the students that required an ability θ from those who do not have it. The higher the value of “a”, the greater the inclination of the Item Characteristic Curve (ICC) and the more discriminant an item is (Zanon *et al.*, 2016).

As a result, most tests used in item response theory consist of multiple-choice items. Therefore, the answers in the first phase were scored dichotomously: the correct answer receives a score of one and an incorrect answer generates a score of zero. From the data transformation, we estimated the 2PL model using the mirt R package (Chalmers *et al.*, 2012) from R 3.6.3.

### 4.2 Domain proficiency

The number of correct answers in each of the three test domains (Bioinformatics, Biology, and Computer Sciences) was assessed for each team, and different proficiency levels were determined by the percentage of correct answers in each domain. The proficiency levels analyzed in each area of knowledge were 80%, 70%, 60%, and 50% of correct answers. This metric is useful for analyzing the heterogeneity of members’ profiles of each team. We also analyzed overlaps in proficiency in different domains, and ranking shifts between teams when comparing the overall ranking with the rankings of each test domain.

### 4.3 Multivariate analysis

Each question in the second phase of LBB was annotated with its respective number according to the order presented in the exam with a letter indicating the order of the sub-question. The 30 teams with scores greater than zero were grouped into three clusters groups of ten. Teams were named 1 to 30 and sorted by the final score. We labelled the groups into three divisions: (i) Group 1: 1th-10th; Group 2: 11th - 20th and Group 3: 21th-30th. Then, a PCA-biplot was performed using the ggbiplot v0.55 package in R 3.6.3.

## 5 Outcomes

### 5.1 First phase

The first phase of the LBB 2019 comprised 59 teams and 168 competitors. The summary of demographic information showed the participation of 64% men (cisgender/transgender), and 31% women (cisgender/transgender) (Figure 2-A). The participants were majority students, accounting 37% of undergraduate, 35% of master, and 21% of Ph.D.. (Figure 2-B). All competitors were distributed majority in Biological Sciences (49%) and Exact and Earth Sciences (31%)(Figure 2-C). Furthermore, competitors were individually assessed on Bioinformatics areas of knowledge (Figure 2-D), Programming languages (Figure 2-E), and level of programming (Figure 2-F). From a total of 127 answers, 34% stated to have more knowledge in genomics, followed by software development (around 31%) and transcriptomics (around 30.7%). Furthermore, more than 50% of the competitors declared using Python as a programming language, followed by R and Perl. The first phase reached 18 out of 27 Brazil’s federative units (67%) (Figure 2-G), demonstrating the LBB coverage across the country.

**Fig. 2.**
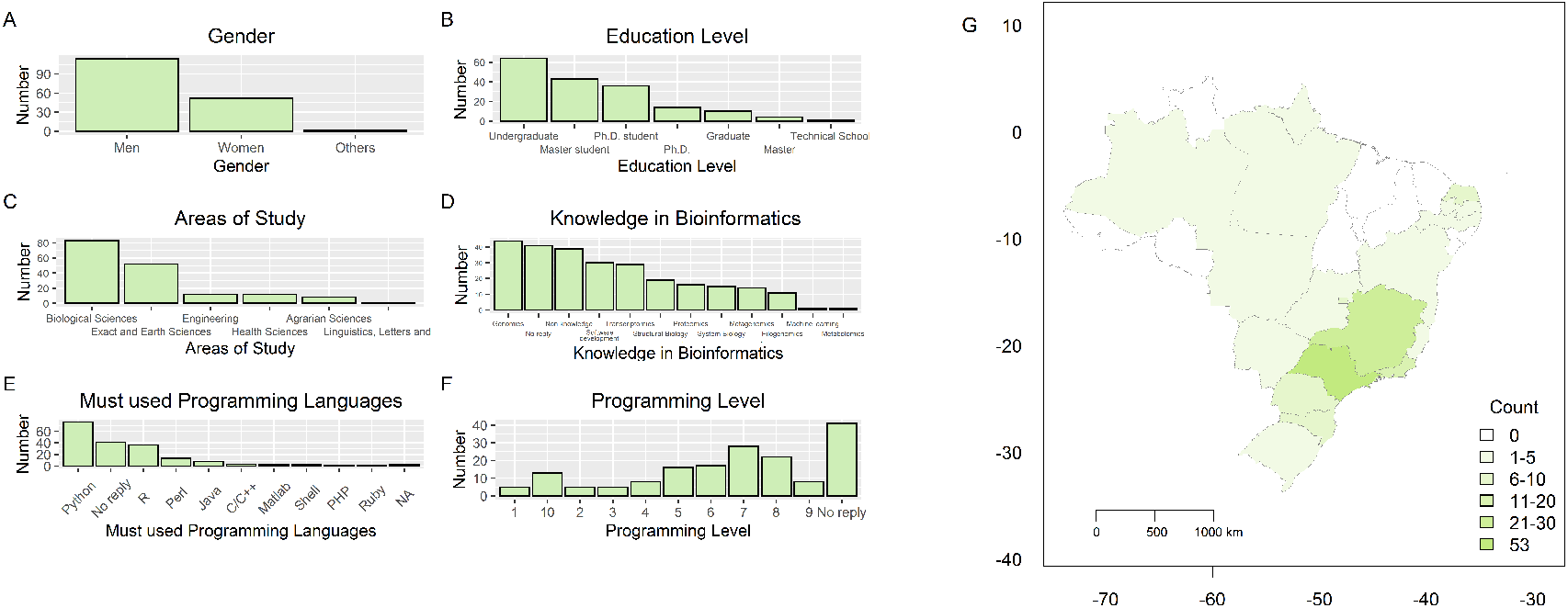
LBB competitors profile. The summary of the demographic information showed: A) gender, B) educational level, C) areas of study, D) previous knowledge in Bioinformatics, E) programming language used, F) programming level, and G) distribution of participants’ universities of the LBB 2019 by affiliation state.

Out of the 59 teams, 5 teams (8.5%) no-show in the competition. Considering teams that submitted the test during the 1st phase, we calculated an average of 34.89 / 60 possible points, and a median of 35/60 points, with the lowest score reaching 4 points, and the highest, 48. If we take into account the score by area of knowledge, the average score was 12.17 for Computer Science, 12.46 for Biology, and 10.65 for Bioinformatics. The competition rule stated that the teams should reach at least 50% of the test to move on to the second phase, which means that the threshold was 30 points. In our analysis, we also observed that if the criteria were 50% in each area of knowledge, we would have a decrease of 25% (11 teams of 44) on the approved teams for the second phase.

During the first phase we observed a total of 22 multiple-choice questions with less than 50% correct response rate, which included seven, six and nine questions from Computer Science, Biology and Bioinformatics, respectively.

Considering the score obtained in each question, we calculated the Item Characteristic Curve (ICC) and Item Information Curve (IIC) (Figure 3). The graph demonstrated the importance of each item in the discrimination identified by the model. The information curve for each item and its ability, as well as its contribution to the model, generated from the ICC, can be found in Figure 4 (Table S1 contains the values “a” and “b” found by the model 2PL test). Also, the characteristic curve of each item in the first phase of LBB, as well as its contribution to differentiate the teams, are shown in Figure 5, in which the difficult items are shifted to the right of the scale, indicating the higher ability of the respondents who endorse it correctly.

**Fig. 3.**
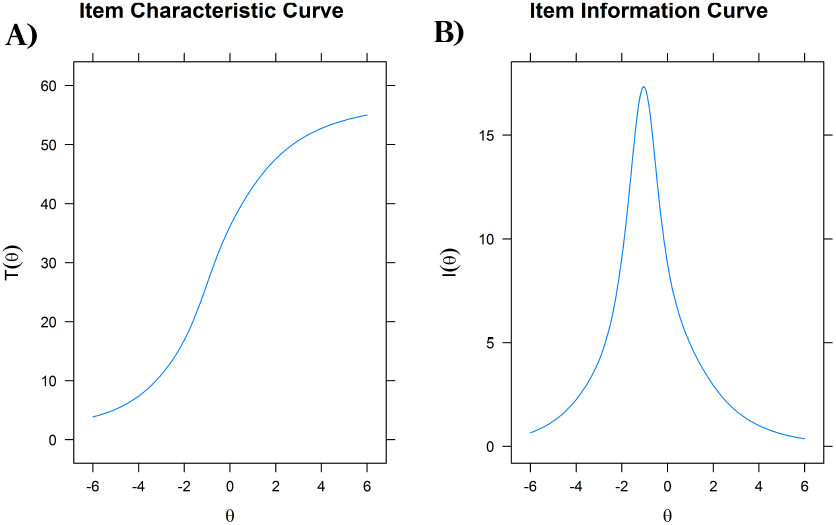
(A) Item Characteristic Curve (ICC) for the responses of the teams participating in the first phase of the LBB. The x-axis represents the team’s ability (θ) to achieve a score (T(θ)). Due to the characteristic of the model’s curve, we can determine that the first phase test was able to separate participants with low and high ability. (B) Item Information Curve (IIC) of the 2PL model. The IIC for the whole test shows that the test provides the most information for slightly-higher-than average ability levels (about θ--1.5), and provides much information about extremely high or low ability levels.

**Fig. 4.**
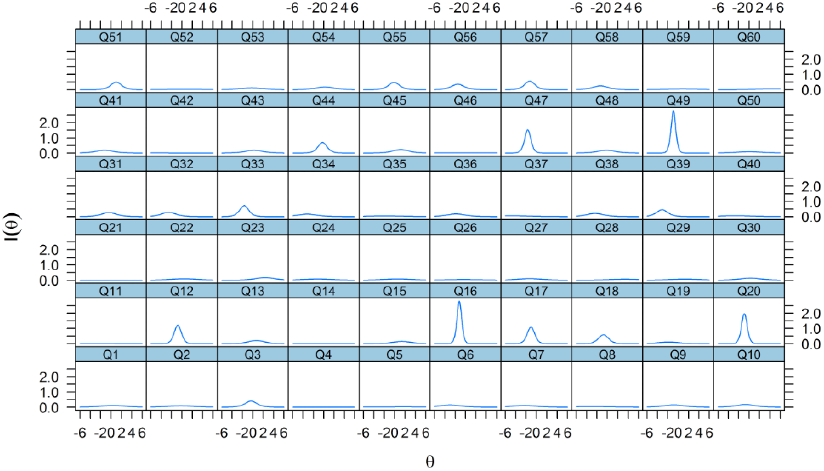
Item Information Curve (IIC) for all questions from the first phase of the competition. The x-axis represents the ability θ of each participant and the y-axis the information of a participant with the ability θ aggregate to the model (I(θ)). The greater the information provided by the item, the greater it aggregates to the model.

**Fig. 5.**
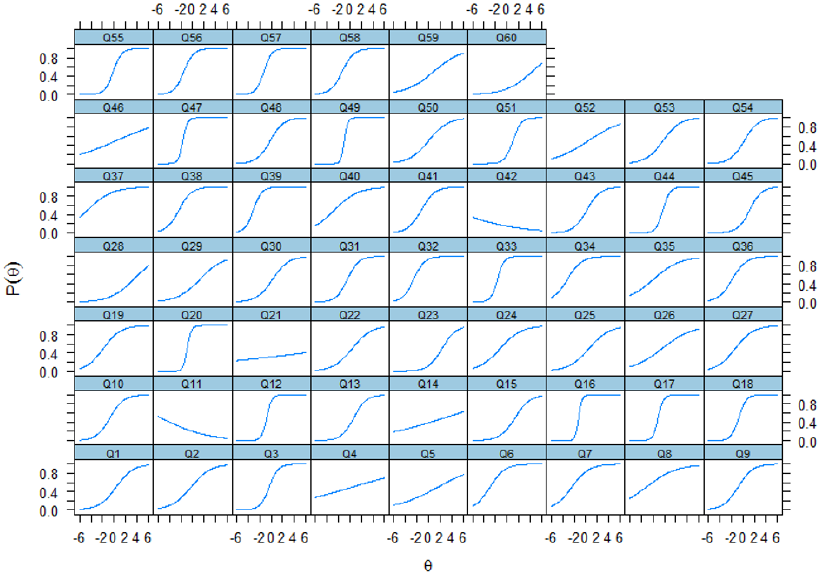
Item Characteristic Curve (ICC) for all questions from the first phase of the LBB. The x-axis represents the ability *θ* of each participant and the y-axis the probability of a participant with ability θ getting the item right (P(θ)). The greater the ability required, the greater is the probability of getting the question right

In Figure 4, the implication is that the higher the discrimination parameter, the greater the information provided by the question. Each question is associated with a specific discrimination factor according to its hit rate. It implicates that each question will have a different graph associated. The questions that are contributing the most with information to the model are the Q49 of Bioinformatics and Q16 of Computer Science.

This result means that participants who had a low score, missed these questions, but those who had a high score got them right. The questions with low ability level (low values of θ) contributed more to the model, and it’s confirmed analyzing the Item Information Curve (IIC) of the model (Figure 3-B). We can also see in Figure 5 that the slope of the curve of these mentioned questions is very pronounced, indicating greater importance in the division of the participants with greater and lesser ability.

When we analyzed the top 10 questions with the greatest ability to distinguish students (“a” values in Table S1), we detected 5 questions in Computer Science (Q16 (highest value), Q20, Q12, Q17, Q18), 4 questions in Bioinformatics (Q49, Q47, Q44, Q57), and just one question in Biology (Q33). This demonstrates that the subjects that will most distinguish the teams were Computer Science and Bioinformatics. Therefore, teams with greater knowledge of these areas had a greater ability and achieved a higher score. It may be explained by the fact that only around one-third of the participants came from a computer science-related background. In addition, when asked about the participants’ programming level, although the distribution has an average of 7 when told to list all programming languages that they can work. Script-based languages were the vast majority of participants chosen (Python or R), middle-level languages, as C or C++, were cited just once. It raises an insight into the lack of computer science knowledge in bioinformatics formation in Brazil.

Analyzing the 10 most difficult questions (top 10 “b” values) in Table S1, we noticed that there were four biology questions (Q21 (highest value), Q28, Q23, and Q29), three computer science questions (Q14, Q5, and Q15) and three bioinformatics tests (Q60, Q45, and Q51). This result indicates that there was a homogenization of difficulties throughout the competition. The ICC can also highlight problematic questions as can be seen in Q11, a tricky and specific question about the usage of coevolutionary constraints in protein sequence annotation and structural prediction; and Q21, a question about Mendel’s Laws. In the next editions, this methodology can be applied to guide the identification of problematic questions before the release of the results.

Another interesting observation is the fact that the teams did not always exhibit a similar level of proficiency in all test domains (assessed by the number of correct answers in each domain). In fact, when teams are ranked by the number of correct answers in one of the test domains, the ranking of the whole test is never replicated. The correlation between the test ranking and the ranking of the domain bioinformatics is 0.8662, indicating that Bioinformatics was the domain that best represented the performance of teams in the test. The ranking of computer science correlates to the general ranking with a correlation of 0.8293, and Biology showed the worst correlation, with a value of 0.7722. As shown in Figure 6, this reflects the fact that many well-performing teams in Biology had poor overall scores, on the other hand, poor performing teams in Biology showed good overall scores.

**Fig. 6.**
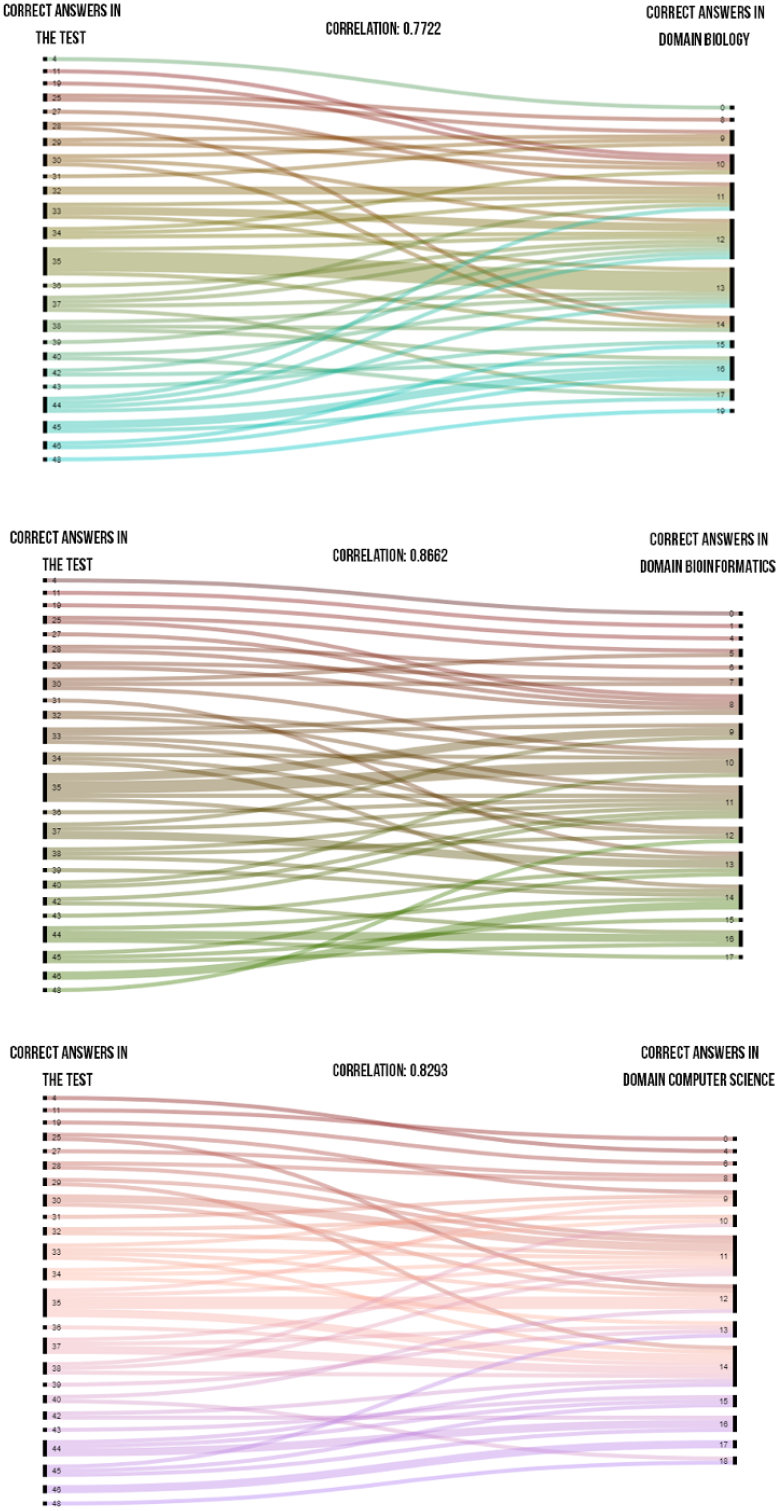
Scores correspondences analysis by domain. The first column represents the overall score in the test, and the second column the score in the tests for each domain. The associated correlation between each score is also represented.

When different levels of proficiency were assessed in each test domain, it was possible to observe that proficiency in Bioinformatics is the rarest proficiency among teams, independent of cutoff used to determine proficiency (Figure 7-A). Also, the most common overlap in proficiencies occurred between Biology and Computer Science (Figure 7-A), reinforcing the importance of our effort to promote multidisciplinary teams in the LBB-Match and other social media interactions (Figure 7-B).

**Fig. 7.**
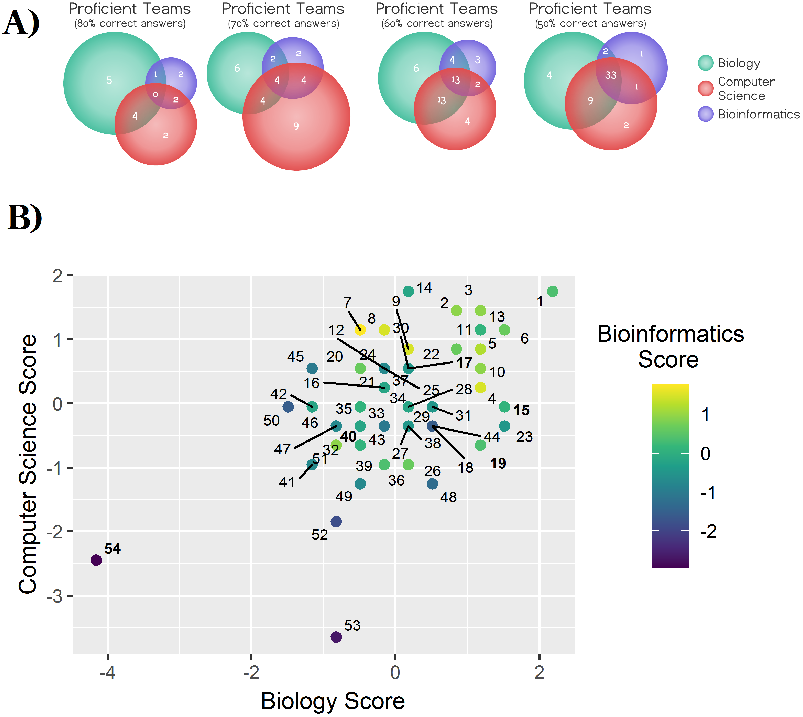
(A) Venn diagrams of domain proficiency, measured by percentage of correct answers in each domain and (B) Scatter Plot of normalized scores for Computer Science (Score.CS) and Biology (Score.Bio). The dot points are colored according to the Bioinformatics score (Score.Bioinfo). The groups highlighted in bold are from LBB Match.

To verify if teams differ in relation to their bioinformatics scores, as observed in the proficiency discrimination analysis, we performed a scatterplot of the normalized scores of Computer Science and Biology, with the color intensity equal to the normalized score in Bioinformatics. In addition, the teams were labelled by the final position in the first phase of LBB. We observed that there were no higher scores in Bioinformatics in the teams that were better positioned, thus showing that the bioinformatics proficiency was variable. Figure 7-B still concludes that many teams received the average score in both Computer Science and Biology (indexed around the coordinate (0, 0)).

### 5.2 Second phase

In the second phase, there were 44 participating teams. The participants had three days to complete five computational biology challenges totalling 50 points. The questions were implemented on the Stepik platform. The challenges needed to be deterministic problems, computationally solvable or at least able to infer the optimal scenarios. The challenges could have distinct evaluation equations, but deterministic questions were scored according to the correct answer and the non-deterministic questions were evaluated according to the distance from the optimal result. Considering all teams’ scores, the average reached 25.61 points and the median reached 31.27 points.

We conducted a biplot (Figure 8), which includes the position of each question in terms of PC1 and PC2, and also shows how the questions are mapped into it (plot of loadings).

**Fig. 8.**
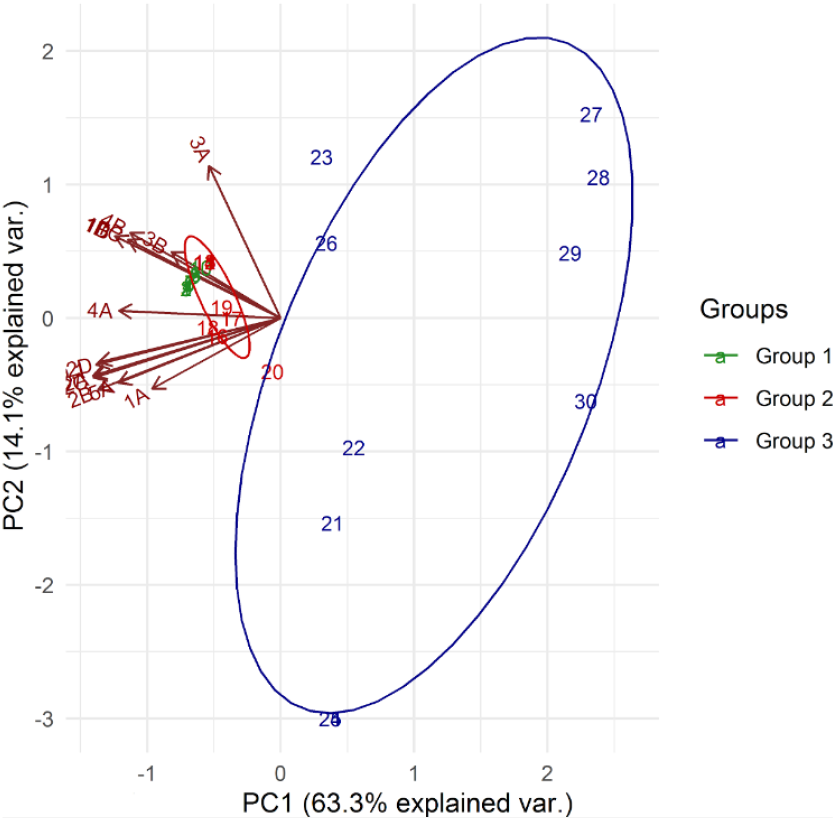
PCA biplot from the scores of the 30 groups that obtained a non-zero score in LBB second phase. Legend: Group 1: top10 teams with a high score; Grupo 2: teams in 11th to 20th position ranked by the final score. Grupo 3: 21th to 30th position ranked by the final score.

The higher score groups were characterized by high values for 1B, 1C, 1D, 3A, 3B, and 4B, meaning these questions were important to split teams with high and low scores. Moreover, the teams with lower scores clearly grouped in a distinct cluster overall the plot with the negative importance of loadings.

### 5.3 Third phase

Upon reaching the third phase, the top-three teams had to submit a research project and received scores for each criterion judged. Through data available from public databases, the finalist teams developed a research project including the scientific question, justificative, objective, methodology, preliminary results and schedule. A fictitious financial funding was set at R$ 200,000.00 (two hundred thousand reais) and the time to conclude the project was set in 1 (one) year.

The evaluation of each project included the extent reached, the training of human resources, the social return, the environmental impact, as well as the creation and support of new collaboration networks, infrastructure, dissemination of results and among others. The teams prepared a research project in a maximum of five pages to be handed over to the organizing committee one month before the end of the competition.

The final test was an oral presentation with a minimum of 15 minutes and a maximum of 20 minutes. Afterwards, an oral argument was carried out with a maximum duration of 40 minutes.

To maintain transparency and facilitate the assessment methodology, the organization has set up an assessment form in which the total score will be 65 points, with the possibility of increasing the score to up to 90 points (bonus score). In the event of a tie, the judges will be summoned to a meeting, where, in a vote, they will issue the final ranking.This form was made available to everyone on the website, encouraging non-finalists to read the questions and when evaluating the presentation they could think about the project.

All questions and criteria used are set out in the EVALUATION FORM in the supplementary material.

Figure 9 shows the average score of the three finalist teams by type of question addressed (Scientific Question, Choice of Methodology, Discussion and Results, and Presentation). With this analysis, we observed the criterion with the highest score, and those that obtained the most variation (standard deviation value). Among the criteria considered, “Presentation” and “Choice of Methodology” were the criteria with the highest and lowest scores, respectively. This shows that, although the team’s presentation was good, the methodology chosen in the project presented deficiencies, which was totally expected considering that our main audience were students. The “Discussion and Results” and “Presentation” criteria obtained the largest standard deviations, showing that these criteria impacted more than all others in the result. Question 9 (Q9) was the question with the highest standard deviation, in which it verifies whether the results presented positive social return and environmental impact, showing that it was not considered by the finalists teams. Also, the referees had an option to grant the teams up to 5 extra points for outstanding performances in any aspect not considered in these analyzes, which was important to define the results.

**Fig. 9.**
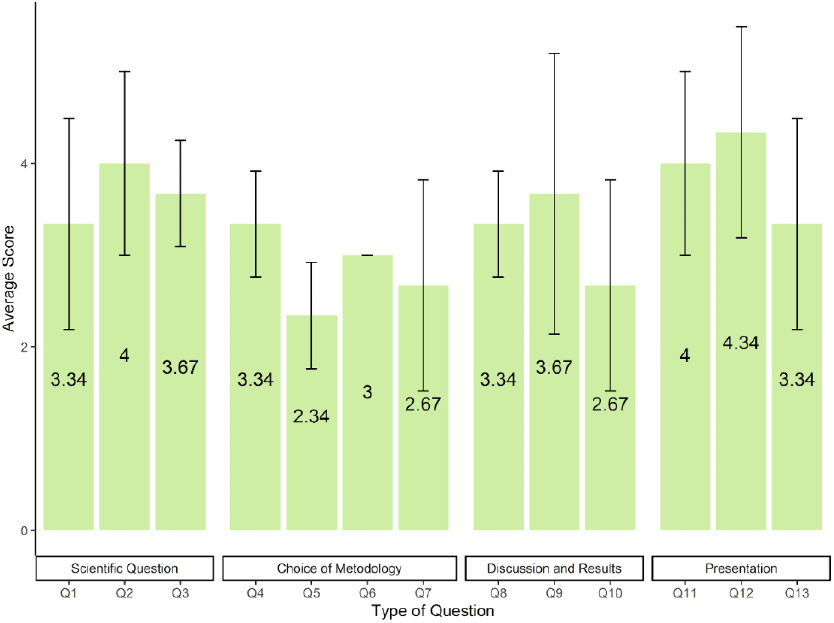
Outcomes of the third phase. The average score of the three teams in each of the 13 questions by each criterion. All questions are set up in EVALUATION FORM in the supplementary material.

## 6 Discussion

Nowadays, Bioinformatics in Brazil is a cutting-edge field with an open road of possibilities and developments, in several different sectors using bioeconomy, and genomic-based solutions. Thus, teaching capacities and delivery transfer knowledge are still only demanded by universities and research centers working with Bioinformatics and Computational Biology. Overcome this problem, we organized the League of Brazil Bioinformatics (LBB), the first competition in Computational Biology in Latin America. It was a wonderful opportunity to access and test the capacities of new talents in the country. It was designed and assembled by a majority of graduate students, motivated by the continuous training in Bioinformatics.

The LBB 2019 reached more than 160 people at different levels of education, programming skills and areas of study (Figure 2). This demonstrates that greater integration of the bioinformatics community was promoted, in addition to providing training for people at different levels of study, and fomenting professional development. In addition, the magnitude of this competition reached competitors in 67% of Brazilian states (Figure 3), demonstrating that integrative initiatives in the Brazilian bioinformatics society can further increase this percentage and reach states that Bioinformatics is not yet introduced. Furthermore, the coverage of Brazilian states by LBB is more related to regions in which graduate programs in Bioinformatics are already consolidated, such as Minas Gerais (UFMG), São Paulo (USP), Paraná (UFPR and UFTPR), Rio de Janeiro (FIOCRUZ and LNCC) and Natal (UFRN).

In relation to the test performance, the Item Response Theory (IRT) demonstrated an excellent separation between the teams that performed the test in the first phase (Figure 3). The items with the most information added to the model were those of Bioinformatics and Computer science, thus showing that the teams that have more skills in these requirements would do well. However, the skill required by the teams in the biology test remained constant, since only one question contributed effectively to the model.

Our analysis showed that the number of questions and the time was adequate, however, we realized that instead of applying a threshold of 50% to the test, using 50% of each area could select stronger teams, which could increase the learning during the LBB by putting competitors from different backgrounds together. Also, we believe that by imposing a minimum score in each test domain, teams will be encouraged to study and train in more diverse areas of knowledge.

The low participation of biology questions in the IRT model could mean that the questions in this test domain were easier, however, when taking into account the number of participants with a background in biological sciences, it is also very likely that the candidates were, in general, more proficient in this test domain. This was corroborated by the proficiency analysis performed, as well as by the low correlation between the ranking of the teams in the first phase and the ranking of the teams in the biology domain since even some well-performing teams in the first phase performed poorly in Biology.

Bioinformatics was shown to be the most problematic test domain for the competitors, and we believe this reflects the deficiency in a truly multidisciplinary training. This is especially conspicuous when we compare the number of proficient teams in Bioinformatics with the number of proficient teams in both Biology and Computer Science. It is likely that teams were multidisciplinary because the participants in a team had different backgrounds, but the participants themselves didn’t present a multidisciplinary background.

In the second phase, issues of Computational Biology divided the groups with the highest score in relation to those with the lowest scores. It allowed selection of the top-three team, however with a small difference between the top 15 teams. These results highlighted the importance of revamp the second phase of the next LBB edition by increasing the number of questions, changing the difficulty of the challenges and reducing the time.

The third phase demonstrated the team’s ability to create a functional, applicable research project with important criteria in the current bioinformatics scenarios. The evaluation form proved a good start, but it may have included a bias in the final result.

After the analyses of LBB 2019, we decided to revamp the LBB regulation. Our first step was to rearrange our work structure and include new members of the organizational team (Figure 10), allocating them into workgroups to be trained. In the meantime, we changed the regulation to embrace reassessment of issues after the tests, increase the difficulty of the second phase through time reduction (for two days), and increase the number of tests (ten challenges). Also, we started to prepare the LBB 2021 and automate the possible processes to replicate the LBB 2019 with the needed improvements. Furthermore, we are planning the next steps to expand the competition to Latin American countries.

**Fig. 10.**
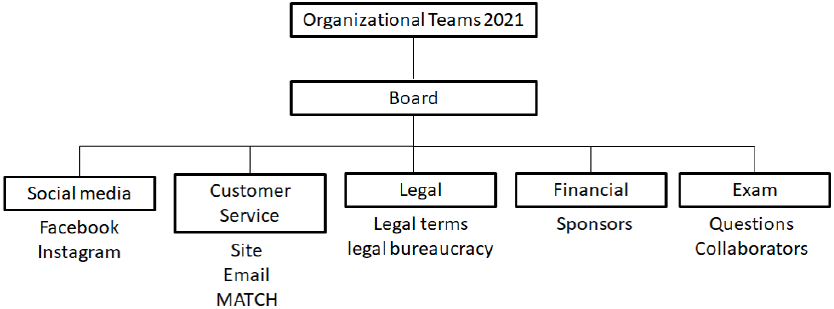
LBB 2021 organization. To organize the project, it was divided into groups to optimize people. The social media group is responsible for disseminating information in instagram and facebook. The customer awareness group is responsible for the oficial channels including email and site, aiming answering questions, and providing instructions for the competition. The legal group takes care of legislation related to both regulation and country legislation. Financially group responsible for fundraising. The Exam development group is responsible for making the questions and checking the questions of the invited professors.

## 7 Conclusion

The League of Brazilian Bioinformatics (LBB) proved to be efficient in bringing together the Brazil Bioinformatics and Computational Biology community, in addition to stimulating, providing knowledge, and integrating a large part of the country’s students. Many participants found the experience extremely interesting since there is a lack of challenges to stimulate development in this area. In addition, the feedback from the LBB Match participants was generally positive, which thanks to the systems they were able to form a team, participate in the competition and meet people. At this time, we can conclude we were able to attend all proposed objectives by the LBB 2019. Due to the great effort to organize and promote the League, our perspective is to carry out the project on a bi-annual basis. In this way, the next League will take place in 2021. Additionally, we believe that the organizing team is putting significant improvements in place for future events.

## Supporting information

League of Brazilian Bioinformatics Regulation - 2019

Supplementary tables and evaluation form

## Acknowledgements

We would like to thank all the participants of the first LBB; the collaborating professors: Dr. João Carlos Setubal (USP), Dr. Marcelo Brandão (UNICAMP), Dr. Raquel Minardi (UFMG), Dr. Tetsu Sakamoto (UFRN), Dr. Alan Mitchell Durham (USP); the board of the third phase: Dr. Ana Carolina Guimaraes (Fiocruz - RJ), Dr. Jonas Gaiarsa (TAU-CG) and Dr. Milton Nishiyama (Butantan Institute); the event sponsor: Hospital Israelita Albert Einstein Varstation (https://varstation.com/pt/sobre-nos/); and the Brazilian Association for Bioinformatics and Computational Biology (AB3C). Also, we would like to extend our gratifications to Sheyla Treflich for developing the LBB logo, Tiago M. Z. M. Gouveia for helping with insights about data analysis of the first phase, and Alice Barros Câmara for collaborating in translating the regulation.

