## Supplementary material for "League of Brazilian Bioinformatics: a competition framework to promote scientific training": League of Brazilian Bioinformatics Regulation - 2019

### REGULATION TO 2019 EDITION

#### 1. General Provisions

1.1. The Brazilian League of Bioinformatics (LBB) is an achievement of the ISCB Regional Student Group Brazil (RSG-Brazil) in partnership with the Brazilian Association of Bioinformatics and Computational Biology (AB3C).

1.2. The planning and execution of LBB and its activities are the responsibility of the Organizing Committee.

1.3. LBB's organizational support structure is located at AB3C's headquarters at Rua do Matão, 1010, Cidade Universitária, São Paulo / SP, CEP: 05508-090.

1.4. LBB is a competition aimed mainly at students from Brazilian universities and the Bioinformatics community, excluding professors and researchers from universities and federal institutes.

1.5. The LBB website is https://lbb.ime.usp.br.

#### 2. Goals

2.1 LBB's main objectives are:

2.1.1. Stimulate the continuous training of human resources in Bioinformatics through participation in competitions.

2.1.2. Train human resources for the production of events in bioinformatics.

2.1.3. Stimulate and promote the organisation of future Bioinformatics competitions, both nationally and internationally.

2.1.4. Promote the integration of the Bioinformatics community in the country and encourage collaboration between LBB participants.

#### 3. Participation requirements

3.1. Teams will be admitted to participate in LBB, under the following conditions:

3.1.1. All participants must be over 18 at the time of registration.

3.1.2. The teams must be composed of 2 or 3 members.

3.1.3. The teams might be composed of members with different educational levels. At least one member has to be enrolled in a higher education institution.

3.1.4. People from any area and academic backgrounds are accepted.

3.1.5. Only 1 (one) member with a complete doctorate per team will be admitted at the registration time (up to two years of degree).

3.1.6. The teams might be composed of members from different institutions.

3.2. It is not allowed registration of professors and researchers hired in universities or federal and state institutes as team members.

3.3. Members of the Organizational Committee, operational coordinators, and collaborators professors are not allowed to register in LBB.

#### 4. Registration

4.1. Registration must be performed by one team member (called the leader).

4.2. The group leader must present proof of enrollment in a Brazilian higher education institution.

4.3. Only one form per team will be accepted. In the case of duplicate registration, only the last form will be considered.

4.4 Each member can only enroll in one team at the LBB.

4.5 The registration form will be available on the LBB website.

4.6 Team registrations will be accepted from 8 AM (BRT) on May 1st until 23:59 (BRT) on June 30th.

4.7 LBB will not be responsible for applications not received due to technical issues and network congestion, therefore it is recommended to send applications in advance.

#### 5. Duties of the Organizing Committee and LBB Coordinators

5.1. LBB is composed by the organizing committee, operational coordinators, and partner teachers.

5.1.1. Organization Committee

5.1.1.1. Responsible for organising the LBB and appointing the board to prepare and correct the questions and challenges.

5.1.1.2. At least one member of the organising committee will be nominated by the president of RSG-Brazil.

5.1.2. Operational coordinators

5.1.2.1. They will be appointed by the organising committee, as demanded by it.

5.1.3. Partner professors or postdocs

5.1.3.1. They will be required to elaborate questions for the first and second phases, according to the needs evaluated by the Organizing Committee.

5.1.3.2. They will be responsible for evaluating the projects developed in the third phase of LBB. The evaluation will happen during the X-meeting 2019.

5.1.3.3 They will be appointed by the Organizing Committee.

5.1.3.4 It is a prerequisite to have a Ph.D.

#### 6. Evidence structure and correction criteria

6.1. The LBB will be carried out in 3 phases:

6.1.1. First phase

6.1.1.1. The first phase will begin on August $4^{th}$ at 1 PM (BRT), with a classificatory and eliminatory character.

6.1.1.2. The exam will last 5 hours and 3 minutes (five hours and three minutes).

6.1.1.3. The first phase will consist of 60 (sixty) multiple-choice questions equally divided into three areas: Biology, Computer Science, and Bioinformatics.

6.1.1.4. Each question will be worth 1 (one) point in the first phase grade.

6.1.1.5. Teams with scores below 50% in the exam will be eliminated from the LBB.

6.1.1.6. The answer form must be submitted once. In the case of sending more than one form per group, only the first submission will be considered.

6.1.1.7. LBB will not be responsible for not receiving exams as a result of any technical problems and network congestion, so it is recommended to send the test in advance.

6.1.2. Second phase

6.1.2.1. The second phase will be carried out in 3 (three) days, from September $13^{th}$ to $15^{th}$ with a classificatory and eliminatory character. The test will be available from 8 AM (BRT) on September $13^{th}$ and will be available until 11:59 PM (BRT) on September $15^{th}$.

6.1.2.2. The second phase will consist of 5 computational biology challenges.

6.1.2.3. The challenges of computational biology will be automatically corrected, and the specific score of each challenge will be detailed in the statement. The expected resolution to the challenges will be exact or approximate, depending on the challenge.

6.1.2.4. Only the submitted output is going to be analysed.

6.1.2.5. Each challenge will be worth 10 (ten) points in the final score.

6.1.2.6. Only the three teams with the highest scores will be accepted in the third phase of the LBB. The result will be made available within one week after the closure of resources.

6.1.2.7. In case of a tie, the following tiebreaker criteria will be used, in the following order:

6.1.2.7.1. Score obtained in the first phase.

6.1.2.7.2 The lesser number of questions that scored zero in the second phase.

6.1.2.7.3. Submission time in the first phase.

6.1.2.8 After the previous scenarios, if the tie remains, all teams will do an extra challenge with the date to be published by the Organizing Committee.

6.1.2.9 LBB will not be responsible for exams not received as a result of any technical problems and network congestion, so it is recommended to send the test in advance.

6.1.3. Third phase

6.1.3.1. The third phase consists of the development of a project outlined by the Organizing Committee. The final results of the projects must be presented by the teams at the X-meeting 2019.

6.1.3.2. If any team member can be in-person at X-meeting 2019, the team must present its results in the form of a webinar.

6.1.3.3. LBB will not be responsible for problems in the presentations or webinars of the finalist teams as a result of any technical problems. Therefore, the submission of the presentation files must be done with the advance stipulated by the Organizing Committee.

6.1.3.4. Project presentations must be made in the format of 15-minute seminars, explaining the scientific question and the methodologies used, as well as the results obtained.

6.1.3.5. The evaluation criteria include:

6.1.3.5.1. The existence of a well-defined scientific question.

6.1.3.5.2. The appropriate choice of methodologies to answer that question.

6.1.3.5.3. The exploration and proper interpretation of the results obtained.

6.1.3.5.4. The clarity and creativity of the project presentation.

6.1.3.6. The judging panel will be responsible for evaluating the participating groups and nominating a winning team.

6.1.3.7. The winner will be announced during the X-meeting 2019.

6.2. The dates for all phases will be defined in the official calendar and will be published on the LBB webpage.

6.3 The results of the three phases, as well as the final score of all teams during the competition, will be published on the official website of the event.

6.4 We will not allow contestation of the results for any LBB phases.

**7. Place of the tests**

7.1. The first and second phase tests will be carried out online on a platform published by the Organization Committee on the official LBB website.

7.2. The third phase will take place online or during the X-meeting 2019 conference, Campos do Jordão – SP, Brazil.

7.3. The tests will be available only for the date and time stipulated in the schedule. The teams must submit their answers through the indicated platform.

7.4. LBB is not responsible for technical failures in the transmission of responses to the examining board and the candidate is entirely responsible for ensuring that responses are sent.

**8. Awards**

8.1. The award ceremony will be held during the last day of X-Meeting 2019.

8.2. All team members from the finalists' teams will receive fee exemption to X-meeting 2019 registration under the AB3C rules.

8.3. The prizes will be announced on the official LBB website.

8.4. The certificates of all participants and awarded students will be made available in digital format and will be sent by email.

**9. Final considerations**

9.1. Any omissions in this regulation will be analyzed and decided in a sovereign and unappealable manner by the Organizing Committee of the Brazilian League of Bioinformatics.
