## Supplementary tables and evaluation form for "League of Brazilian Bioinformatics: a competition framework to promote scientific training"

**Table S1**. Values of a (discrimination) and b (difficulty) of the 2PL model of each knowledge area for the 60 questions in the first phase of the LBB.

| Knowledge Area | Question (Qi) | a | b |
|---|---|---|---|
| CS | Q1 | 0.677184614 | 0.375594022 |
| CS | Q2 | 0.595669378 | -0.2544022 |
| CS | Q3 | 1.267648793 | -0.35025751 |
| CS | Q4 | 0.150213833 | 0.497767792 |
| CS | Q5 | 0.272514146 | 1.691122277 |
| CS | Q6 | 0.77492097 | -2.96670947 |
| CS | Q7 | 0.6710295 | -2.39132228 |
| CS | Q8 | 0.378753189 | -3.1230163 |
| CS | Q9 | 0.788256406 | -0.85130745 |
| CS | Q10 | 0.849993797 | -0.69126061 |

| | | | |
|---|---|---|---|
| CS | Q11 | -0.248475874 | -5.55864787 |
| CS | Q12 | 2.215946279 | -0.79666983 |
| CS | Q13 | 0.923859688 | 0.683360721 |
| CS | Q14 | 0.160472304 | 2.835483322 |
| CS | Q15 | 0.772714504 | 1.389237722 |
| CS | Q16 | 3.341538465 | -1.21066181 |
| CS | Q17 | 2.094052667 | -1.05596877 |
| CS | Q18 | 1.534820666 | -0.73431154 |
| CS | Q19 | 0.667034087 | -1.86253556 |
| CS | Q20 | 2.806777303 | -0.92381105 |
| B | Q21 | 0.068107356 | 11.43157047 |
| B | Q22 | 0.606629371 | 0.409721461 |
| B | Q23 | 0.828012131 | 2.363200184 |
| B | Q24 | 0.544660247 | -1.18185507 |
| B | Q25 | 0.557793044 | 0.582622738 |
| B | Q26 | 0.387371707 | -0.38903123 |
| B | Q27 | 0.630909939 | -1.32078153 |
| B | Q28 | 0.494180906 | 3.409392409 |
| B | Q29 | 0.51348152 | 1.266124635 |

| | | | |
|---|---|---|---|
| B | Q30 | 0.723545627 | 0.243147962 |
| B | Q31 | 1.015831514 | -0.41560245 |
| B | Q32 | 1.047449654 | -2.56363653 |
| B | Q33 | 1.706633895 | -1.6695737 |
| B | Q34 | 0.793477913 | -3.19110215 |
| B | Q35 | 0.428787816 | -1.88049401 |
| B | Q36 | 0.825573677 | -1.84544957 |
| B | Q37 | 0.471296434 | -4.59676435 |
| B | Q38 | 0.908781745 | -2.38646555 |
| B | Q39 | 1.338924916 | -3.11217277 |
| B | Q40 | 0.490613358 | -2.67551154 |
| BioInfo | Q41 | 0.864003856 | -1.24198323 |
| BioInfo | Q42 | -0.184877153 | -9.52685142 |
| BioInfo | Q43 | 0.855765405 | 0.216642178 |
| BioInfo | Q44 | 1.670270262 | -0.2286126 |
| BioInfo | Q45 | 0.969547183 | 1.166217418 |
| BioInfo | Q46 | 0.219858468 | 0.343448598 |
| BioInfo | Q47 | 2.469140236 | -1.68054068 |
| BioInfo | Q48 | 0.863796224 | -0.17682688 |

| | | | |
|---|---|---|---|
| BioInfo | Q49 | 3.339488506 | -0.94045481 |
| BioInfo | Q50 | 0.602862149 | 0.013697615 |
| BioInfo | Q51 | 1.400070285 | 0.998039735 |
| BioInfo | Q52 | 0.329866431 | 0.467024694 |
| BioInfo | Q53 | 0.67605638 | -0.22453786 |
| BioInfo | Q54 | 0.835970161 | 0.220037391 |
| BioInfo | Q55 | 1.373215187 | -0.11766919 |
| BioInfo | Q56 | 1.182776545 | -1.51483846 |
| BioInfo | Q57 | 1.496766107 | -1.28644172 |
| BioInfo | Q58 | 1.00167053 | -1.32871098 |
| BioInfo | Q59 | 0.42158735 | 0.932015018 |
| BioInfo | Q60 | 0.451975647 | 4.368745145 |

Legend: CC: Computer Science; B: Biology; BioInfo: Bioinformatics

EVALUATION FORM

1. Scientific Question

Criterion 1: The research challenges are perfectly formulated and placed against the state of the art and the existing literature.

| **0** | **1** | **2** | **3** | **4** | **5** |
|---|---|---|---|---|---|

Criterion 2: Original, bioethical, and internationally competitive research project.

| **0** | **1** | **2** | **3** | **4** | **5** |
|---|---|---|---|---|---|

Criterion 3: Innovative research project.

| **0** | **1** | **2** | **3** | **4** | **5** |
|---|---|---|---|---|---|

2. Choice of Methodology

Criterion 4: The methodology described is convincing.

| **0** | **1** | **2** | **3** | **4** | **5** |
|---|---|---|---|---|---|

Criterion 5: The research project can be carried out within the proposed deadline.

| **0** | **1** | **2** | **3** | **4** | **5** |
|---|---|---|---|---|---|

Criterion 6: The research project is reproducible.

| **0** | **1** | **2** | **3** | **4** | **5** |
|---|---|---|---|---|---|

Criterion 7: The project is feasible according to the budget.

| **0** | **1** | **2** | **3** | **4** | **5** |
|---|---|---|---|---|---|

3. Discussion and Results

Criterion 8: The results have a great possibility of significantly expanding the frontier of knowledge in the area and, therefore, of having a very relevant scientific impact.

| 0 | 1 | 2 | 3 | 4 | 5 |
|---|---|---|---|---|---|

Criterion 9: The results have a positive social return and environmental impact.

| 0 | 1 | 2 | 3 | 4 | 5 |
|---|---|---|---|---|---|

Criterion 10: There is the creation/support of new collaboration networks, infrastructure, dissemination of results, and among others.

| 0 | 1 | 2 | 3 | 4 | 5 |
|---|---|---|---|---|---|

4. Presentation

Criterion 11: Clean and creative presentation.

| 0 | 1 | 2 | 3 | 4 | 5 |
|---|---|---|---|---|---|

Criterion 12: Oral presentation of the team.

| 0 | 1 | 2 | 3 | 4 | 5 |
|---|---|---|---|---|---|

Criterion 13: The writing of the document.

| 0 | 1 | 2 | 3 | 4 | 5 |
|---|---|---|---|---|---|

5. Bonus

Bonus Criterion: The team presented relevant topics beyond the expected.

What was the Bonus? Bonus score of 1-5

Bonus 1:

______________________________________________________________
